# Floral organs act as environmental filters and interact with pollinators to structure the yellow monkeyflower (*Mimulus guttatus*) floral microbiome

**DOI:** 10.1101/721647

**Authors:** María Rebolleda Gómez, Tia-Lynn Ashman

**Affiliations:** Department of Biological Sciences, University of Pittsburgh, Pittsburgh PA 15260

**Keywords:** Microbiome, Pollination, Dispersal, Community assembly, Serpentine seeps, Metacommunity, *Mimulus guttatus*, Anthosphere

## Abstract

Assembly of microbial communities is the result of neutral and selective processes. However, the relative importance of these processes is still debated. Microbial communities of flowers, in particular, have gained recent attention because of their potential impact to plant fitness and plant-pollinator interactions. However, the role of selection and dispersal in the assembly of these communities remains poorly understood. We evaluated the role of pollinator-mediated dispersal on the contribution of neutral and selective processes in the assembly of floral microbiomes of the yellow monkeyflower (*Mimulus guttatus*). We sampled floral organs from flowers in the presence and absence of pollinators within five different serpentine seeps in CA and obtained 16S amplicon data on the epiphytic bacterial communities. Consistent with strong micro-environment selection within flowers we observed significant differences in community composition across floral organs and only a small effect of geographic distance. Pollinator exposure affected the contribution of environmental selection and depended on the rate and “intimacy” of interactions with flower visitors. This study provides evidence of the importance of dispersal and within-flower heterogeneity in shaping epiphytic bacterial communities of flowers, and highlights the complex interplay between pollinator behavior, environmental selection and additional abiotic factors in shaping the epiphytic bacterial communities of flowers.

## Introduction

Community assembly is the product of neutral and selective processes (Nemergut et al. 2013; Vellend 2016). In particular, the composition of a community can change through speciation, dispersal, ecological drift (or sampling of individuals and species over time), and environmental selection (Vellend 2016). Environmental selection is a deterministic process and depends on fitness differences between populations (Chesson 2000; Chase & Leibold 2003; Vellend 2016). Neutral processes, in contrast, are independent of niche differences between species and are predicted to be driven by stochastic differences in birth and death (Vellend 2016). Neutral processes can lead to rapid differentiation of communities when dispersal between them is low (McArthur & Wilson 1963; Hubell 2001; Economo & Keitt 2008). Dispersal, in turn, can be deterministic or stochastic depending on species differences in dispersal abilities (Nemergut et al. 2013; Lowe & McPeek, 2014; Evans et al. 2016). The relative importance of neutral and selective processes in community assembly is still subject of much debate (e.g., Hubell 2001; Tilman 2004; Leibold & McPeek 2006; Morrison-Whittle & Goodard 2015) and the contribution of dispersal to these processes can be hard to measure in the field (Evans et al. 2016). Understanding the relative contributions of these processes in host-associated microbiomes is an important first step to understanding the consequences of microbe communities for the host (Costello et al. 2012).

Studies of host associated microbiomes have highlighted the importance of selection by the host in shaping its associated microbial communities (e.g., Rawls et al. 2006; Ofek-Lalzar et al. 2014; Pratte et al. 2018). The host can favor colonization and growth of certain microbes over other through diverse mechanisms like: immune system activity (e.g., Donaldson et al. 2018), host secretions (e.g., Schluter & Foster 2012; Ofek-Lalzar et al. 2014), or specific environmental characteristics like high osmolarity in flower nectar (Herrera et al. 2010). These effects, however, can be overpowered by dispersal from other hosts or the host environment (Burns et al. 2017). Thus, understanding the relative contributions of drift (i.e., neutral) and selective processes in the host can provide insight on the drivers of host-associated microbiome assembly, their changes over time (e.g. Burns et al. 2016), and the potential sources of these microbes (e.g. Venkatamaran et al. 2015). Recently, flower-associated microbiomes have been established as an excellent system to study community assembly and metacommunity dynamics (Belisle et al. 2012; Shade et al. 2013, Vannette & Fukami 2017; Chappell & Fukami 2018).

Flowers are multi-purpose reproductive structures and microbial communities of flowers can have a large impact on plant fitness by directly affecting the survival and reproduction of the plant (e.g., Alexander & Antonovics 1995), or through effects on pollination (Vannette et al. 2013; Herrera et al. 2013; Schaeffer et al. 2014; Schaeffer et al. 2017; Rering et al. 2017). Understanding microbial community assembly in flowers, can highlight important, and underappreciated, ecological processes affecting floral evolution and plant-pollinator interactions.

Despite significant variation across floral organs, and potential effects of microbes of anthers, pollen, styles and stigma on direct fitness effects (not mediated by pollinators), most studies of microbial communities associated with flowers have concerned microbes of nectar (e.g., Herrera et al. 2010; Belisle et al. 2012; Pozo et al. 2016; Mittelbach et al. 2016; Vannette and Fukami 2017). These studies have shown the importance of pollinators in shaping some of the assembly patterns of these microbiomes (Belisle et al. 2012; Herrera et al. 2013; Vannette and Fukami 2017). But there is substantial variation in microbial composition that is not explained by the presence/absence of pollinators (e.g. Vannette and Fukami 2017), and the source of most floral microbes remains unknown. Each floral organ is likely to create unique conditions for the establishment of bacteria (Aleklett et al. 2014; Junker & Keller 2015). Pollinators that transport microbes have different behaviors on flowers and could create varying opportunities for contact with floral structures (Laverty & Plowright 1988; Russell et al. 2019). Yet we have a poor understanding of the extent to which floral organs and their interaction with pollinators creates heterogeneity in microbial communities within flowers.

In this paper we address the relative importance of neutral (i.e., drift and passive dispersal) and selective processes (i.e., organs that create unique habitats within the flower), as well as their interaction with pollinator-mediated dispersal in shaping epiphytic bacterial communities in a flower with no nectar production. If neutral effects are the main factor explaining community assembly, then we expect that: different organs will have a random phylogenetic representation of the whole flower metacommunity and that the most abundant microbes in the whole metacommuity will also be the most frequent in the different organ samples. In addition, if communities geographically farther apart are less likely to share microbial migrants, then, under a neutral model, we expect that these distant communities will be more likely to diverge as a result of ecological drift (Hubell 2001; Soininen et al. 2007). Thus, we would expect that spatial location of the plant in the habitat and not floral organ to explain most of the differences between communities, and that community differentiation (beta diversity) will increase with geographical distance. In contrast, if the different floral organs act as selective microenvironments, then we expect that floral organ and not plant geographic location will be the main determinant of community composition. In addition, if pollinators are the major agents of microbial dispersal, then we would expect pollinator exclusion to affect differentiation among locations or organs. Specifically, pollinators could homogenize communities by transporting microbes across large spatial scales, or they could increase differentiation by moving microbes mainly within local patches. Finally, high rates of pollinator-mediated dispersal could overwhelm the effects of local dynamics of environmental selection within the flower.

## Materials and Methods

### Study system and species

The yellow monkeyflower, *Mimulus guttatus* (*Erythranthe guttata*, Phyrmaceae) is self-compatible, hermaphroditic and insect-pollinated annual/perennial herbaceous plant that is a dominant component of the serpentine seep communities in northern California (Harrison et al. 2000; Freestone & Inouye 2006; Arceo-Gómez & Ashman 2011). Flowers are zygomorphic and tubular, produce little or no nectar, and there is high variability among flowers in the quality of the pollen rewards (Robertson et al. 1999; Wu et al 2008). In the field, monkeyflowers interact with a variety of insect pollinators of varying behaviors and sizes (Arceo-Gomez & Ashman 2014; Koski et al. 2015). As a result, seep monkeyflower is highly generalized and well-connected within the pollinator networks of these serpentine seeps (Koski et al. 2015).

Monkeyflowers were studied within five seep communities at the McLaughlin Natural Reserve in northern California, USA (Table S1). These seeps are characterized by a high diversity of flowering species restricted by abiotic factors (e.g., water availability) and are separated in space by a grassland matrix. Therefore, seeps can act as discrete replicate communities (Harrison et al. 2000; Arceo-Gómez and Ashman 2014).

### Experimental set up and epiphytic bacterial sampling

At the height of flowering in 2017 (May 9-20) we established five transects along five serpentine seeps with three (RH1, RHA and RHB) or four (TP9 and BNS) sampling points (Table S1; Fig. 1A). Within each transect the locations of the sampling points provided a range of inter-seep distances from ~10m to ~100m (Fig. 1B). The geographical position of the longest sampling point within a seep was recorded as GPS coordinates (Table S1). For the shortest distances we used the distances measured in the field. Using qGIS 2.18.10 (qGIS development team, 2016) we projected all coordinates on WGS84/UTM Zone 10N to obtain a distance matrix in meters.

**Figure 1.**
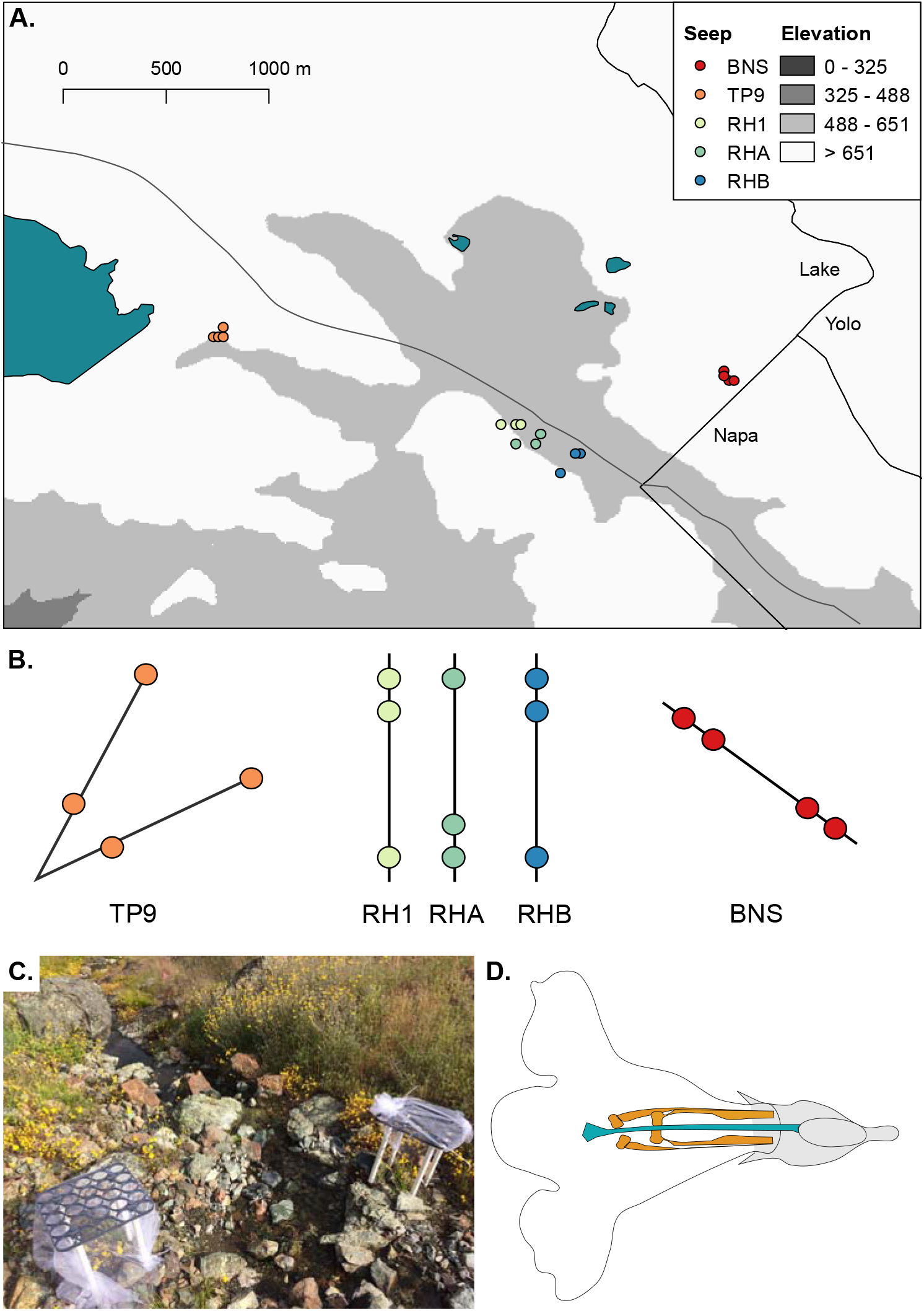
Experimental design. A. Map of seeps at McLaughlin Natural Reserve and sampling locations within them. The different colors indicate each seep. B. Diagram of sampling locations within each seep. Distances between sampling location as 10-100m. C. Picture of pollinator exclusion cage and control cage in one of the locations. D. Diagram of *Mimulus gutattus* flower showing the floral organs studied: petals (pink), the four stamens (yellow) and the style (turquoise).

Within each seep and at each location, we set up paired control and pollinator-exclusion cages (treatments). Cages were constructed from PVC and tulle (Joann Fabrics, ITEM #1102979). The control cages had open sides to allow for visitation (Fig. 1C). Wearing sterile gloves, we marked the petiole of several flower buds per plant within a cage with a permanent marker. After marked flowers were open for 3-4 days we carefully dissected the organs of three flowers from two to three different plants using sterile forceps. The stamens (anthers and filaments), petals (only the corolla, without the calyx) and styles with stigma (no ovary) were stored in separate sterile vials (Fig. 1D). To obtain enough DNA, for each sampling location-treatment combination we pooled replicate organs from three different flowers, ending up with one sample for each organ, for each treatment and location (102 samples of floral organs).

To get a better idea of the potential sources of floral microbes at each location we also sampled a basal aerial leaf from each monkeyflower plant, soil (one random location per seep) and flowers from the co-flowering community (39 samples total). The community samples were a mix of five flowers representing the species in the local community (2 x 3m plot; Table S1). All samples were collected at the same time (within a week of each other) to minimize changes in co-flowering community, pollinator community and other environmental variables like temperature.

### DNA extraction and 16S rRNA amplicon sequencing

To obtain the epiphytic bacterial communities of the flower organs (or leaves or flower communities) we washed samples with 1ml of sterile phosphate buffered saline (PBS) and vortexed them for ten minutes to detach bacterial cells from the tissue (in previous tests we did not observe differences in colony forming units between five minutes of sonication with a small jewelry sonicator and ten minutes of vigorous vortexing; data not shown). We concentrated the microbial cells and used only the bottom 250μl of the pellet for DNA extraction (avoiding the floral tissue). For our soil data we used 200 g of soil directly in our DNA extraction protocol. We extracted DNA from all samples using DNeasy PowerSoil Kit (Quiagen). We added a control for DNA extraction and sampling in the field (maintained a tube with 250μl of sterile water opened in the field for ten minutes). Both of our controls failed to amplify. We sequenced the 16S *rRNA* gene V4 hypervariable region using one run in the Illumina MiSeq platform (Illumina, CA, USA). We used the 515FB-806RB primer pair (FWD:GTGYCAGCMGCCGCGGTAA; REV:GGACTACNVGGGTWTCTAAT) and use paired-end sequencing of 150 base pair per read (Caporaso et al. 2012). All of the sequencing procedures including the library preparation, were performed at the Argonne National Laboratory (Lemont, IL, USA) following the Earth Microbiome Project protocol (http://www.earthmicrobiome.org/protocols-and-standards/16s/) with 12-bp barcodes. To reduce chloroplast contamination, we used peptide nucleic acid (PNA) clamps (Lundberg et al. 2013). All sequences will be made available in NCBI’s Short Read Archive.

### Sequence processing

We used PEAR v0.9.10 paired-end merging (Zhang et al. 2014). After sequencing we obtained a total of reads and we were able to successfully pair (92%). After merging our reads, we re-assigned the barcodes to the merged reads with a custom script written by Daniel Smith (https://www.dropbox.com/s/hk33ovypzmev938/fastq-barcode.pl?dl=1”). Subsequently, we demultiplexed our samples and removed low quality reads (Phred quality scores < 20), aligned them with PyNAST (Caporaso et al. 2010), and assigned taxonomy (OTUs at 97% similarity) with the RDP classifier (Wang et al., 2007), using the Greengenes database (13_8 release), as implemented in QIIME v.1.9.1 (Caporaso et al. 2010b). We removed mitochondria and chloroplast sequences as well as OTUs in low abundance (often spurious) across samples (>0.0005% mean abundance) according to recommendations based on simulations (Bokulich et al. 2012).

OTUs are a conservative measure of variation that clusters together sequences within 97% similarity. Sequencing errors often fall within that 3% of variation that is allowed within the same OTU and are, thus, not interpreted as meaningful biological variation. However, OTU clustering also losses much of the finer biological variation. DADA2 is a model-based approach for correcting amplicon errors while maintaining sequence level variation (i.e., amplicon single variants or ASVs; Callahan et al., 2016). To evaluate the robustness of our results, we also processed our data following the DADA2 pipeline. After assessing the quality of our reads, we trimmed the last ten base-pairs of our reverse reads but left our forward reads untouched to be able to merge them. Most reads (or 94%) were kept after quality filtering and trimming.

After merging reads (we were able to merge 1.2 x 10^7^ of the filtered and trimmed reads, 92%), we removed chimeras and sequences either too short (less than 248bp) or too long (more than 256bp), and then, we removed chloroplast and mitochondrial sequences. Overall, results from the QIIME 1.9 pipeline (OTUs) and DADA2 (ASVs) were consistent. However, DADA2’s finer resolution tended to amplify stochastic variation between individual samples (see results and discussion). Thus, in this paper we present the results from our OTU (QIIME 1.9 pipeline) data and discuss when both analyses are discordant. Both pipelines are available in gitHub (github.com/mrebolleda/OrganFilters_MimulusMicrobiome).

### Pollination observations

To evaluate the effects of pollination intensity on the microbiome assembly, at each sampling location we observed a similar number of flowers within a 2 x 3m plot and recorded floral visitors for fifteen minutes. A visit was recorded as contact with a monkeyflower or a ‘community’ flower in our plot (we did not count as separate visit multiple visits by the same insect to the same flower, and we only counted the first three visits of the same insect, thus we scored visitation at the plot level). We scored visits to focal monkeyflowers as ‘external’ when a visitor contacted petals or ‘internal’ when an insect contacted anthers or stigma within the flower. We classified each visit by the functional group of the insect following Koski et al. (2015). Each sampling location was observed twice between 9:20 and 12:00, and twice between 12:00 and 16:00. For each location we calculated the visit rate to focal monkeyflowers and to the community of flowering plants in each plot (visits/plot/hour).

### Analyses of species composition

Due to evolutionary divergence in chloroplast sequences (Fitzpatrick et al. 2018), despite using the PNA clamps ~80% of our reads were chloroplast and mitochondria sequences and were removed from analysis resulting in samples with anywhere from 211 to 179975 reads and a median coverage of 5986 reads per sample (Fig. S1A-B). To facilitate accurate comparisons across samples, we subsampled each to an even sequencing depth of 1200 reads. This number was chosen as a balance between the depth of sampling within each bacterial community and the number of communities (Fig. S1C-D). To minimize the potential effects of stochastic variation due to low coverage, we obtained an average sample from each of our communities after 10,000 subsamples with replacement. In addition, we performed all of our analyses with data normalized through a variance-stabilizing transformation as implemented in “DESeq2” (Anders & Huber 2010; Love et al. 2014). We obtained the same overall results with both analyses (data not shown), however we present those using our rarefied community for diversity analyses because the results were slightly more conservative. All statistical analyses were performed in R (3.4.4; R Core Team 2017).

To characterize the epiphytic microbial communities of the monkeyflowers we calculated four distance matrices using “Sørensen” (only richness), “Bray-Curtis” (relative abundance), “Unweighted Unifrac” (relative abundance and phylogenetic distance) and “Weighted Unifrac” (relative abundance and phylogenetic distance) (Lozupone & Knight 2005). These indices of beta diversity allowed us to evaluate the robustness of our results as well as the importance of relative abundances and phylogenetic information in our flower samples. We performed a PERMANOVA to evaluate the effects of floral organ, pollinator treatment and seep for each of the community distance matrices. We used the full model including as factors: floral organ, seep, pollinator and the two-way interactions. We evaluated assumptions of homogeneity of group dispersion using the function *betadisper* (Anderson 2006). Only Bray-Curtis showed heterogeneity in the dispersion across organ samples.

In addition, we evaluated the degree of phylogenetic clustering or overdispersion of communities (from a particular floral organ and pollinator treatment) as the deviation from expected phylogenetic diversity. To calculate phylogenetic diversity, we used the total branch length of a given sample (Faith 1992) and the expected phylogenetic diversity was calculated through binomial sampling of the whole metacommunity tree (O’Dwyer et al. 2012). In this case, we defined our metacommunity as all of the monkeyflower samples together. Expected and observed phylogenetic diversities were calculated using the *picante* package (1.7; Kembel et al. 2010).

Next, to evaluate the relationship between geographical distance and community differentiation, we calculated bacterial community composition distance matrices using Bray-Curtis and Unifrac for each floral organ within each pollinator treatment. We then performed Mantel tests on these matrices using the “vegan” package (2.4-6; Oksanen et al. 2018). We adjusted p-values to account for multiple testing using false discovery rate correction (Benjamini & Hochberg 1995). We also assessed the degree of concordance between flower community composition and microbial community composition at each location through Procrustes analysis comparing a non-metric multidimensional scaling (NDMS) of data for each pollinator treatment and organ combination.

Finally, to evaluate the uniqueness of each community we obtained a “core microbiome”. Comparing overlap in “core” taxa only, provides a way to minimize inflation of non-shared taxa by excluding OTUs that might be present only in a few samples. For this analysis we defined the “core microbiome” as the OTUs that were shared by at least 20% of the samples of a given organ/treatment (the maximum cut-off that still provided a large sample of more than a 100 OTUs). In other words, if an OTU was not present in at least 20% of the samples of a given organ in a given treatment then it was not considered part of the core for that organ and in that treatment. With this list of OTUs we calculated overlap across organs.

### Neutral model fit

To determine the potential importance of neutral processes to community assembly, we evaluated the fit of a neutral model for prokaryotic communities (Sloan et al. 2006; Burns et al., 2016). This model is based on the idea that, under neutral assumptions, taxa that are more abundant in the whole metacommunity are more likely to occur in multiple patches (floral samples). Thus, with a single free parameter *m* (that describes the migration rate) the model predicts the relationship between the frequency to which taxa occur in a series of communities (in this case each organ) and their abundance in the whole metacommunty (all of the monkeyflower samples together).

Using the Akaike information criterion, the fit of this neutral model was compared to a null-model (in this case a binomial distribution) that represents random sampling from the “pool community” without drift or dispersal limitations (Sloan et al. 2007; Burns et al. 2016). The 95% confidence intervals around the model were calculated by bootstrapping with 1000 samples. OTUs within the 95% CI were considered to fit neutral expectations of distribution in the metacommunity. OTUs outside the confidence intervals were separated in two categories: overrepresented (present in more samples than would be predicted by their mean relative abundance) and underrepresented (present in less samples than would be predicted by their mean relative abundance). The relative proportions of OTUs in these different categories among floral organs where evaluated with chi-square tests of independence with Bonferroni correction for multiple tests.

To determine the extent to which over- or underrepresented taxa are exclusive to each organ (instead of shared across two or more of the floral structures), we generated a null model to determine, given the number of OTUs in each category, how many would we expect to be shared between at least two organs controlling for the number of OTUs (independently their category). We repeated this sampling process 10,000 times to generate a null distribution for comparison with our observed values. To determine the proportion of OTUs shared across our organ samples we used the *get.venn.partitions* function in the “VennDiagram” package (1.6.20; Chen & Boutros 2018).

### Analyses of potential sources of monkeyflower microbial communities

To understand the relation between flower communities with other local communities of microbes, we obtained the differences in OTU composition (beta diversity) between our focal monkeyflower epiphytic microbial communities (for a given organ) and the communities acting as potential sources (i.e., the rest of the floral organs, monkeyflower leaves, co-flowering community or soil). We calculated total beta diversity and the components due to nestedness and turnover (Baselga 2010) between the focal communities and the potential sources for each seep using the R package “betapart” version 1.5.0 (Baselga & Orme 2012). A large contribution of “nestedness” means that differences between communities are mostly due to subsample (species losses) from the more diverse to the less diverse community. Whereas, a large contribution of “turnover” indicates species replacement across communities (Baselga 2010). For each seep we compared an average source community (to minimize variation due to differences in sample sizes of sources) with all of the communities for a given floral organ in a given seep.

To evaluate whether potential sources differed in their compositional distance to our focal communities, and to investigate the contributions of nestedness and turnover to their overall beta diversity, we performed an ANOVA with beta diversity as the response variable, and component (nestedness or turnover), floral organ and source as factors. We did not include pollinator treatment in this analysis because communities from both of our pollinator treatments showed the same patterns (see results).

Beta diversity indicates differences between potential sources and our focal communities, and the decomposition of beta diversity into nestedness and turnover components highlights the ways in which those communities are different. However, to identify the likely sources of our focal communities (and the uncertainty around these calls) we used SourceTracker as implemented in R (version 1.0.1). SourceTracker is a Bayesian approach that models a sink community as a mix of potential sources, allowing for assignment into an unknown source when part of the sink (focal community) is not like any of the sources (Knights et al. 2011). The code for all of the analyses is available in gitHub (github.com/mrebolleda/OrganFilters_MimulusMicrobiome).

## Results

### Floral organs are the main factor explaining variation epiphytic bacteria community composition

PERMANOVA results were fairly consistent across all beta diversity indices: pollination treatment (exclusion/control), seep and floral organ and their two-way interactions explained between 26% to 33% of the total variation in epiphytic bacteria of monkeyflowers (Table 1). In general, our model was slightly better at explaining species composition alone (Sørensen and Unifrac) than abundance (Bray-Curtis and weighted Unifrac). The presence of OTUs (more than their relative abundances) distinguishes between organs (Table 1; Fig. S2). Floral organ was significant in its contribution to community composition across all the different distance metrics (Fig. 2A, Table 1, Fig. S2), and alone explained between 4% and 11% of the variation (Table 1). Seep was marginally significant (α=0.05) in all comparisons but Bray-Curtis (Table 1) and pollinator treatment was not significant, but the interaction between pollinator treatment and organ was marginally significant across indices (explaining ~3% of the variation; Table 1, Fig. 2A). Overall, floral organ was the only factor that was consistently significant, with seep and interactions between factors being marginally significant in half or more of the analyses (Table 1).

**Figure 2.**
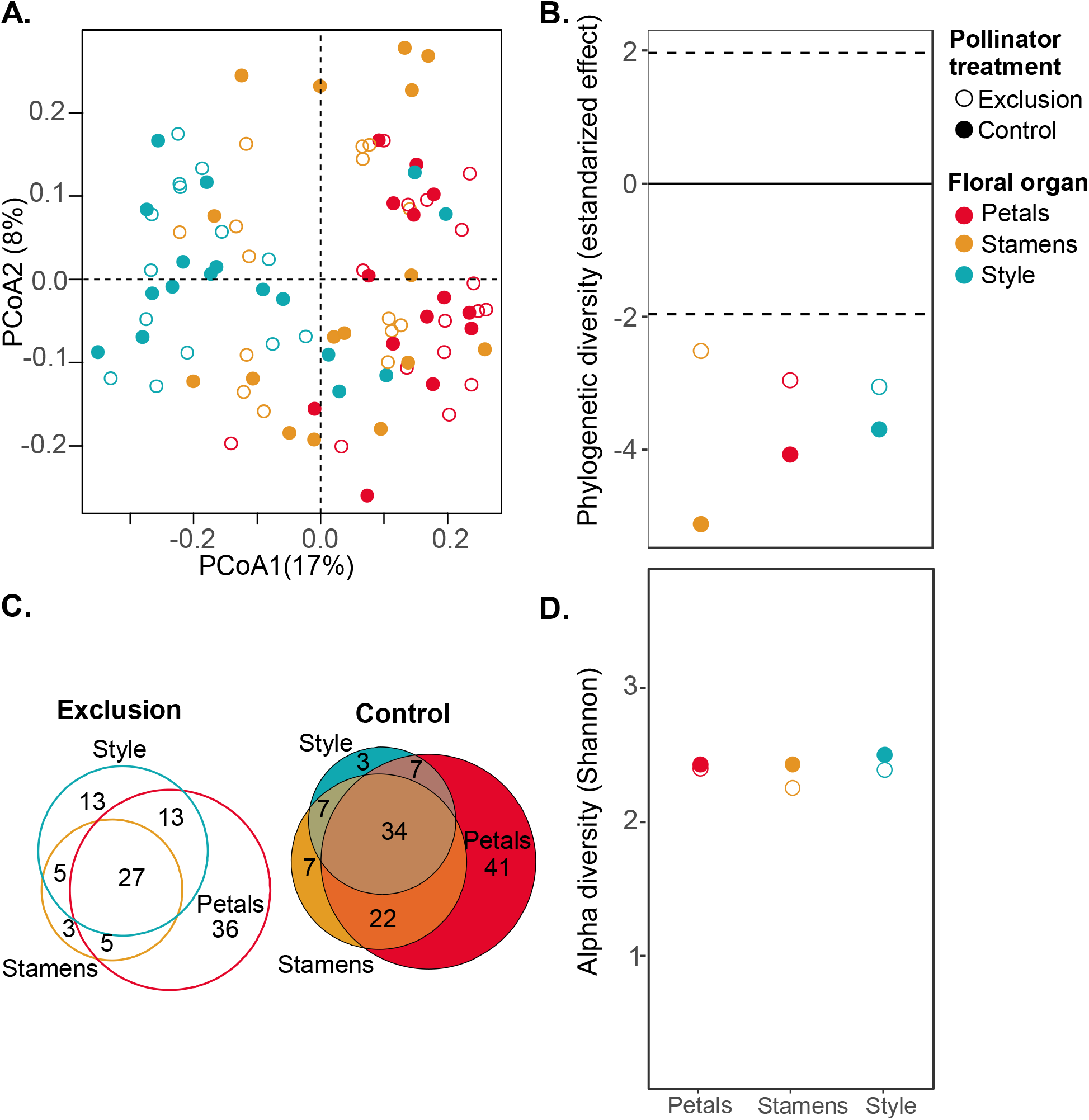
Contribution of floral organ differences to bacterial community structure of monkeyflowers (*Mimulus guttatus*). A. PCoA based on Unifrac differences between samples. Each point represents one sample. B. Bacterial communities of each organ are phylogenetically clustered (i.e., lower phylogenetic diversity than expected). C. Venn diagrams of the core microbiome of each floral organ for each of the pollinator treatments. D. Alpha diversity across floral organs. Small circles represent each sample and the large circles the mean values.

**Table 1.**
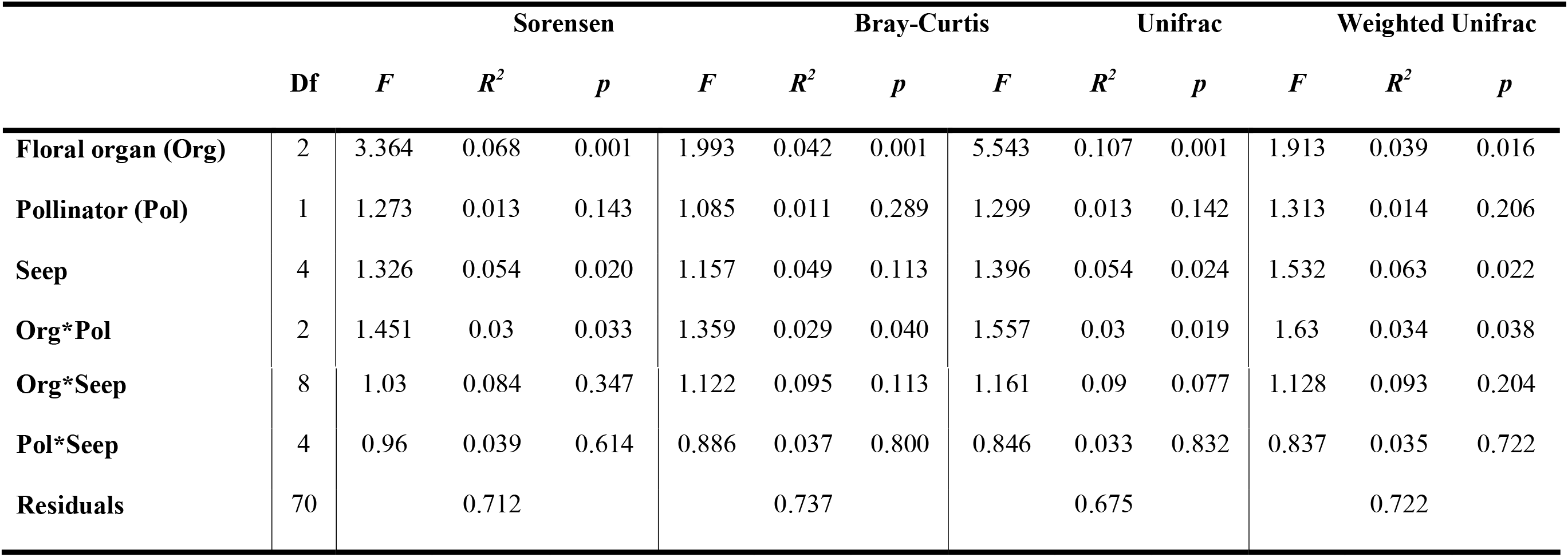
Table with PERMANOVA summaries using different betadiversity indices (Org-Floral organ; Pol-Pollinator treatment).

|  | Sorensen |  |  |  | Bray-Curtis |  |  | Unifrac |  |  | Weighted Unifrac |  |  |
| --- | --- | --- | --- | --- | --- | --- | --- | --- | --- | --- | --- | --- | --- |
|  | Df | F | R<sup>2</sup> | p | F | R<sup>2</sup> | p | F | R<sup>2</sup> | p | F | R<sup>2</sup> | p |
| **Floral organ (Org)** | 2 | 3.364 | 0.068 | 0.001 | 1.993 | 0.042 | 0.001 | 5.543 | 0.107 | 0.001 | 1.913 | 0.039 | 0.016 |
| **Pollinator (Pol)** | 1 | 1.273 | 0.013 | 0.143 | 1.085 | 0.011 | 0.289 | 1.299 | 0.013 | 0.142 | 1.313 | 0.014 | 0.206 |
| **Seep** | 4 | 1.326 | 0.054 | 0.020 | 1.157 | 0.049 | 0.113 | 1.396 | 0.054 | 0.024 | 1.532 | 0.063 | 0.022 |
| **Org\*Pol** | 2 | 1.451 | 0.03 | 0.033 | 1.359 | 0.029 | 0.040 | 1.557 | 0.03 | 0.019 | 1.63 | 0.034 | 0.038 |
| **Org\*Seep** | 8 | 1.03 | 0.084 | 0.347 | 1.122 | 0.095 | 0.113 | 1.161 | 0.09 | 0.077 | 1.128 | 0.093 | 0.204 |
| **Pol\*Seep** | 4 | 0.96 | 0.039 | 0.614 | 0.886 | 0.037 | 0.800 | 0.846 | 0.033 | 0.832 | 0.837 | 0.035 | 0.722 |
| **Residuals** | 70 |  | 0.712 |  |  | 0.737 |  |  | 0.675 |  |  | 0.722 |  |

Higher taxonomic resolution (using ASVs instead of OTUs; see methods) provides additional support for organ as the main factor structuring microbial communities in flowers. Using ASVs we might expect an increase contribution of stochastic and local processes (where specific strains might only be locally distributed, or present in only a few samples). Despite this increased stochasticity, organ is still significant across all diversity indices (Table S2). However, seep and pollinator by organ interaction are no longer significant (Table S2).

Consistent with organs acting as environmental filters, microbiomes of a given organ in a given treatment tended to have less phylogenetic diversity than expected from random sampling of the whole floral bacterial metacommunity (Fig. 2B). Stamens and styles were dominated by OTUs from the Pseudomonadales order, while the most abundant OTUs associated with petals were more evenly distributed across the Pseudomonadales, Bacterioidales and Clostridiales (Fig. S3).

Despite evidence of flower organs contributing to community differences across microbiome samples, we observed high variation across samples of the same organ (Fig. 2A, Fig. S2, Fig. S3) and an exponential decrease in the number of OTUs shared by an increasing percentage of samples of a given organ (Fig. S4A). Furthermore, a large majority of core microbes were shared across organs, with no significant differences between the proportion of shared vs. unique OTUs between pollinator treatments (Fishers exact test, *p*=0.225; Fig. 2C). Among core microbes we found some of the most abundant genus in all organs: *Acinetobacter, Pseudomonas, Bacteroides* and *Corynebacterium*. In addition, we found some common flower associated genus like *Erwinia* and *Lactobacillus* as well as some unidentified *Prevotella* and *Streptococcus* (more commonly associated with human hosts). Across both treatments, petals had the largest number of unique OTUs even though we did not observe differences in alpha diversity at our sampling effort (Fig. 2D).

### Floral organs act as selective filters

We observed no significant relation between geographical distance at the measured spatial scales (meters to kilometers) and beta diversity for petals, and styles, with or without pollinators. Across treatments, communities displayed the same level of differentiation (beta diversity). Only stamens showed increased community differentiation when exposed to pollinators than in the control, and only the stamens exposed to pollinators showed a significant relation of increasing beta diversity with increasing distance (Mantel test with 999 permutations, *r*=0.266, *P*=0.038) (Fig. 3A). This relationship was similar to that seen in the leaf samples (Mantel test with 999 permutations, *r*=0.258, *P*=0.032; Fig 3A).

**Figure 3.**
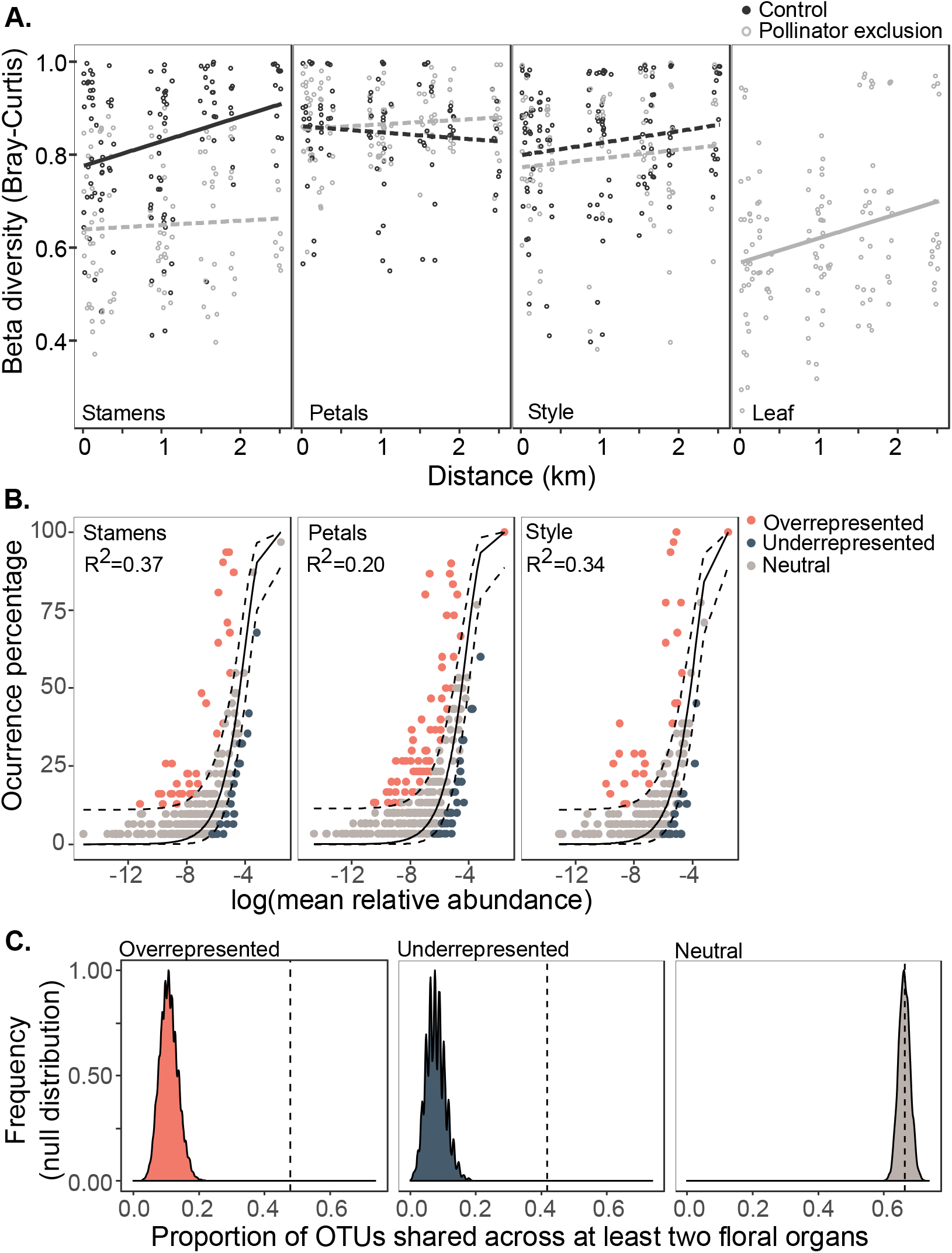
Effect of pollinator treatment in beta diversity across floral organs of *Mimulus guttatus*. A. Beta diversity calculated in a multivariate space from Bray-Curtis distances. B. Relationship between geographical distance and betadiversity (Bray-Curtis) across different floral structures and leaves. C. Proportion of OTUs that are shared across at least two floral organs. The distribution is the result of 1000 simulations assuming no differences between groups other than their sizes. The vertical dashed line indicates the observed values.

Consistent with ecological selection across different floral organs, we found a large proportion of OTUs that are either overrepresented or underrepresented under neutral assumptions (Fig 3B). The neutral model only explains between 20% and 37% of the variation in the distribution of taxa (Fig 3B), but it is a better fit than a model ignoring drift and migration (Table S3). To assess deviations from neutrality across organs, we compared the proportion of OTUs in each category (i.e., overrepresented, underrepresented and neutral) across the three floral organs. We observed significant differences in the proportion of OTUs in each category across the three different organs (*χ*^2^_(4)_ = 19.638, *P*= 0.0006). There was no difference between styles and stamens (*χ*^2^_(4)_= 0.551, *P*= 1), but petals had a higher proportion of overrepresented and underrepresented taxa than styles (*χ*^2^_(2)_ = 15-344, *P*= 0.001) or stamens (*χ*^2^_(2)_ = 11-327, *P*= 0.01). In general, core taxa (present in at least 25% of samples from a given organ) were also some of the most abundant (although this is not always true in the case of some overrepresented taxa that are present in more than 25% of the samples despite having an overall low frequency, and some of the underrepresented taxa, that are not present in even 25% of the samples, but when they are they might be in high abundance; Fig 3B).

The neutral models perform worse when we separate the data by pollinator treatments (i.e. exposed control and pollinator exclusion; Table S3). This reduction could be due to smaller sample sizes. Nevertheless, the neutral models still explain between 15% and 26% in the different treatments of stamens and styles, while these models explain less than 7% of the variation in the distribution of taxa from petals (Table S3). For the most part, we did not observe strong differences in the fit of neutral models when comparing our two pollinator treatments (Table S3, Fig. S4). However, in the presence of pollinators the neutral model explained the data better than in our pollinator exclusion treatment (*R*^2^ Control= 0.15 vs. R^2^ Exclusion= 0.26 and AIC Control= −79.5 vs. AIC Exclusion= −207.3; Table S3).

If each organ is selecting for a particular microbial community, we would expect that over or underrepresented taxa for a particular organ will not be shared across different organs. Whereas OTUs that are distributed according to a neutral model would be distributed more or less randomly across the whole flower. According to our expectations, OTUs distributed according to neutral expectations are shared across two or more organs in the same proportion as we would expect by chance (Fig. 3C). Instead, over and underrepresented OTUs are shared between organs more than we would expect by chance (Fig. 3C). This effect is stronger in the control treatment; in the presence of pollinators over- and underrepresented taxa are more likely to be shared among organs than when pollinators are excluded (Fig. S4).

### Dispersal can overwhelm the effects of organ selection

Despite evidence of organ-specific selection on bacterial communities, we also observed a marginally significant interaction between organ and pollinator treatment (although only in the OTU data). In the pollinator exclusion treatment, almost twice as much of the total variation is explained by floral organ, relative to the control (Fig. 4A). This is true even in the ASV data set, where, organ explains more variation in the pollinator exclusion treatment than in the control (except in the case of Bray-Curtis; Figure S5). In addition, floral organs of pollinator excluded plants have a larger proportion of unique OTUs across a variety of cut-off values for the core microbiome (0-60%; Fig. S4B).

**Figure 4.**
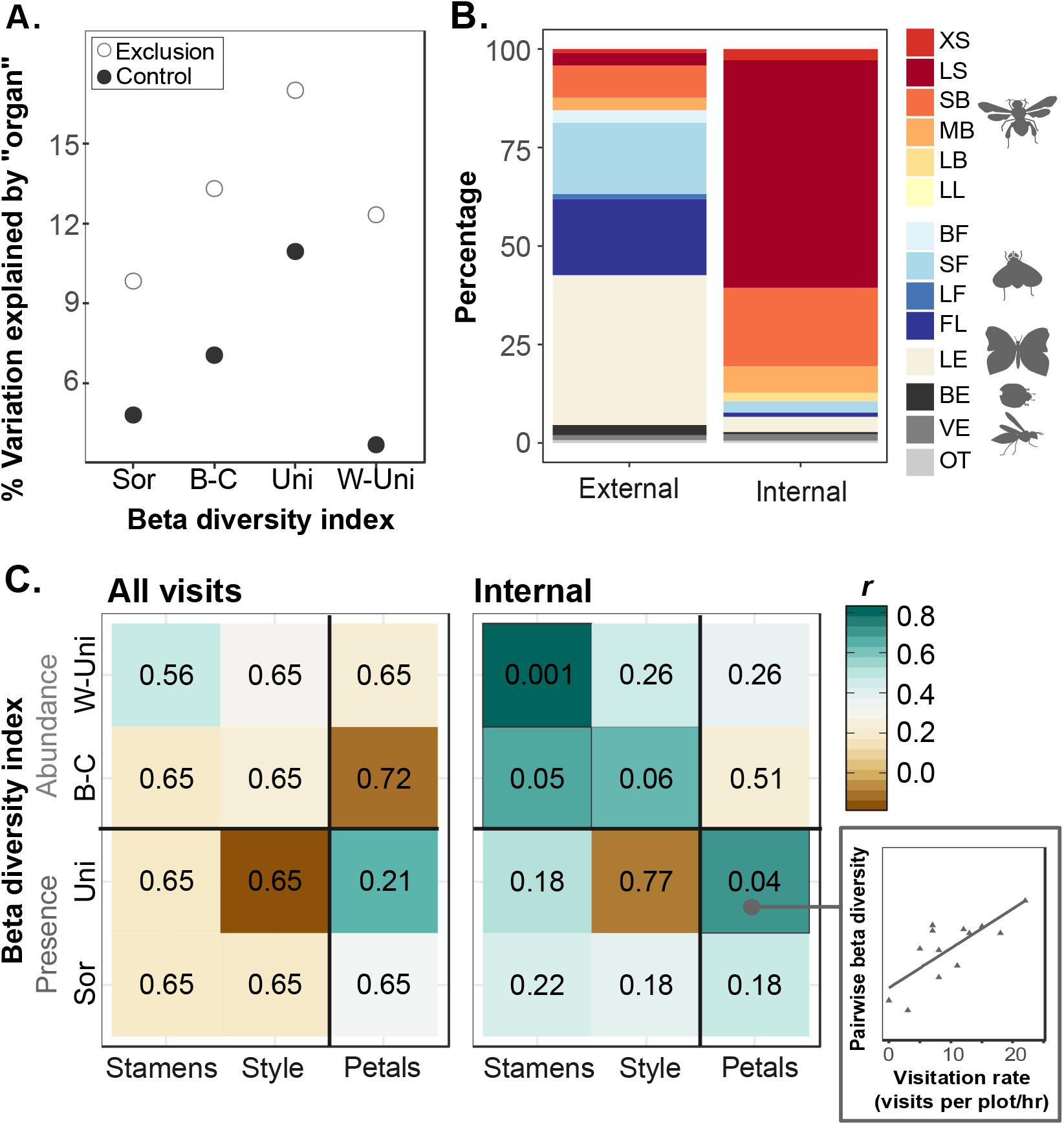
Pollinator effect on community composition. A. Percentage variation in community composition explained by organ in the control and pollinator exclusion treatments. B. Composition of pollinator pool (percentage of total number of visits) contacting external or internal surfaces of the flower (Functional pollinator categories from Koski et al., 2015: XS-extra-large social bees; LS-Large social bees; SB-small solitary bee; MB-medium solitary bee; LB-large solitary bee (pollen on body); LL-large solitary bee (pollen on legs); BF-bee fly; SF-small syrphid fly; LF-large syrphid fly; FL-other flies; LE-moths and butterflies; BE-beetles; VE-wasps; OT-other). C. Correlations between visitation rate to the plots and community composition differences between pollinator treatments at each location (pairwise beta diversity-see small insert). Colors show the strength of the correlation and the numbers inside denote the p-values after adjustment for multiple testing (see text for details). Analyses were performed using different beta diversity indices. Sørensen (Sor) and Unifrac (Uni) include only presence-absence data whereas Bray-Curtis (B-C) and weighted Unifrac (W-Uni) also account for relative abundances.

Pollinator service was heterogeneous in both quality and quantity. Some insects mainly encountered the external parts of *M. guttatus*, whereas others contacted the internal reproductive organs of the flower. The community of ‘external’ visitor insects differed significantly in composition from the community of insects visiting the ‘internal’ parts of the flower (Fishers exact test, *p*>0.0001; Fig 4B). In addition, visitation varied in rate across location with visitation to the yellow monkeyflowers varying from 6-55 mean visits/plot/hour (Table S1). We observed no significant correlations between the total number of visitors (internal and external) and the pairwise beta diversity between petal samples of the pollinator exclusion and the control treatment. In contrast, when we consider only internal visitors (those that might be in direct contact with anthers and stamens) we observed a positive correlation for the stamens (increasing in strength as we account for abundance and phylogenetic information in the beta diversity index used; Fig 4C). This pattern is maintained when looking at the ASV data. In contrast, there are no clear patterns for petals and styles, and all significant correlations (styles using Bray-Curtis and petals using Unifrac; Figure 4C) are lost when analyzing the ASV data (Fig S7).

We did not observe a clear association between the co-flowering community composition and microbial community composition for none of the organ/pollinator treatment combinations (Table S4). Alpha diversity was not correlated with pollinator visitation rates, co-flowering community abundance nor co-flowering community diversity (Fig S8).

### Potential sources of floral microbes

Our results suggest that pollinator-mediated dispersal of microbes can affect community assembly and possibly override the contribution of environmental selection from different floral organs. But the ultimate sources of microbial communities of flowers remain unclear. Using decompositions of betadiversity, we can ask how much organ-specific communities differ from other microbial communities that could act as sources (i.e., soil, *M. guttattus* leaves, heterospecific co-flowering neighbors or remaining parts of the *M. guttattus* flower) and how much of these differences is due to replacement of OTUs (turnover) or loss of OTUs in a nested manner from the potential sources (or more diverse communities). Levels of beta diversity were high across all comparisons (0.77±0.012 SE), indicating differentiation of our focal bacterial communities from all other potential sources. Across floral structures beta diversity was highest when focal organ-specific communities were compared with soil and lowest in comparison with the neighboring heterospecific flowering community. Surprisingly, these patterns of differentiation from potential sources, were not different across pollinator treatments (Fig. 5).

**Figure 5.**
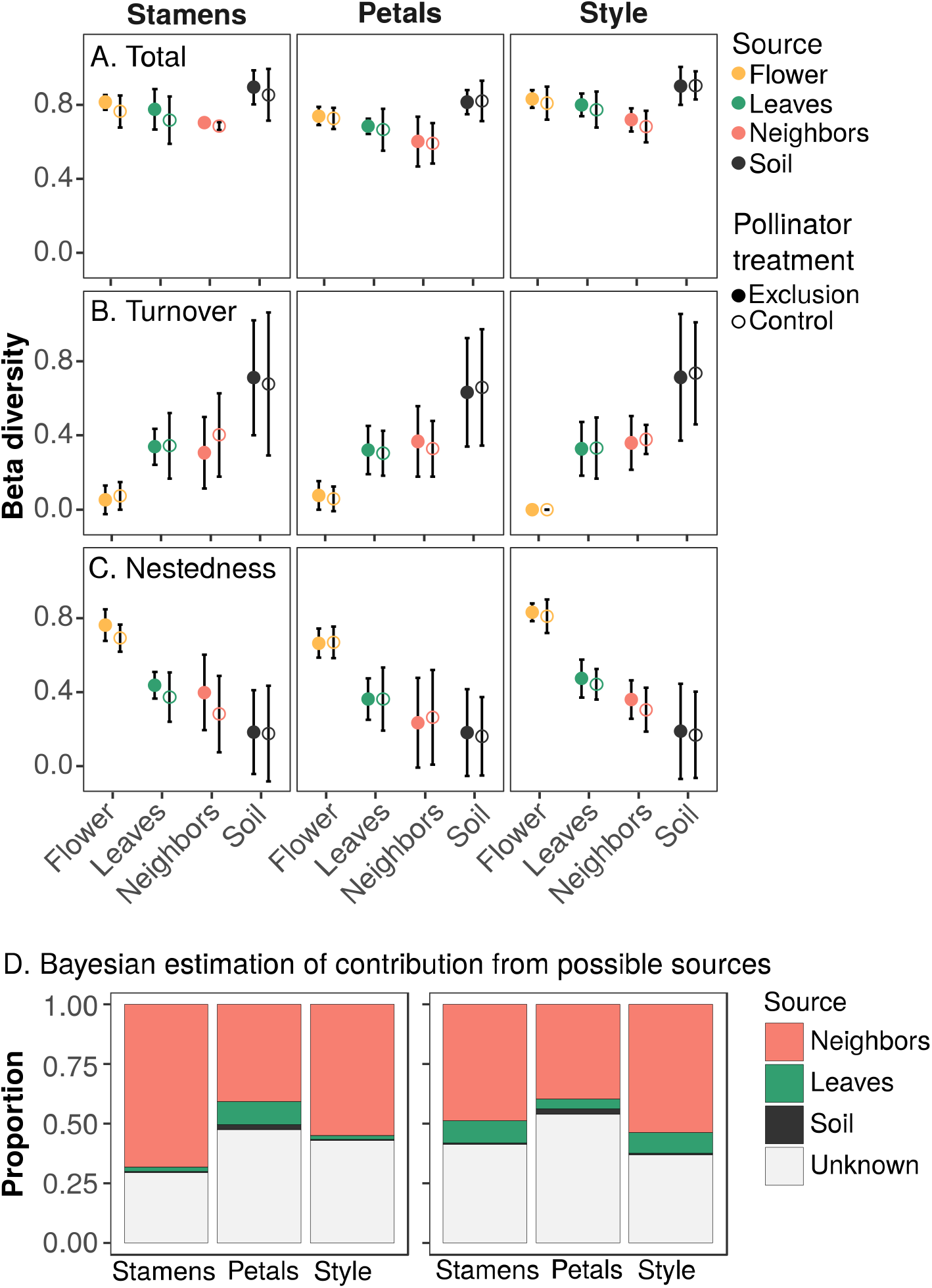
Potential sources of microbial communities in *M. guttatus* flowers. A-C. Components of beta diversity comparing potential source microbial communities (rest of the *M. guttatus* flower, *M. guttatus* leaves, floral neighbors, soil) against the communities of stamens, petals and styles (in columns). A. Total beta diversity as measured by the Sørensen index. B. Portion of beta diversity that is due to turnover of species. C. Component of beta diversity that is due to nestedness. Points are the mean ± one standard deviation. Solid circles are samples from the control treatment and solid circles are from the pollinator exclusion treatment. D. Proportion of OTUs in each treatment that could be assigned to each source using Bayesian estimation. The rest of the flower was not added as a potential source in this analysis.

The contributions of nestedness and turnover were significantly different (ANOVA, *F_(1,84)_*=13.037, *p*= 0.0005) and their contributions varied across sources (ANOVA, *F_(3,84)_*=237.618, *p*<0.0001) but not across floral organs (ANOVA, *F_(2,84)_*=1.758, *p*=0.1787; Fig. 5; Table S3). Comparisons between the focal floral organ and the remaining flower organs had the highest values of nestedness and lowest values of turnover than comparisons with all other potential sources. Soil, instead had the lowest values of nestedness, and the highest turnover (Fig. 5). This analysis suggests that the OTUs in our focal communities are (to some extent) a subset of those present in other flowers in the co-flowering community, and have a number of OTUs not present (or present in low abundance) in our soil samples.

Consistent with these results, using a Bayesian approach to track the potential sources of microbes (Knights et al. 2011), a large proportion of our sample was assigned to be from the co-flowering community (neighbors) and only a small percentage from soil and leaves. All groups (organ and treatment) had a large proportion of bacterial taxa that was assigned to unknown sources, with a larger contribution in the petals.

## Discussion

Despite their importance for plant community function and fitness, we know very little about the communities of microbes that inhabit flowers. Here we showed that different organs within a flower have different epiphytic bacterial communities which overwhelm the effects of geographic distance on community composition. Our results indicate that bacterial communities of flowers are established by the balance of dispersal and environmental selection, but this balance will be different for each organ within a flower. We suggest that floral organs (especially the petals) act as environmental filters and that, in the absence of pollinators, the metacommunity as a whole might be better described within a species-sorting paradigm that emphasizes niche differences (Leibold et al. 2004). However, our data suggest that, within organ environmental selection could become overwhelmed by pollinator-mediated dispersal of new taxa (especially in organs with extensive pollinator engagement like the stamens) and, with high rates of visitation the metacommunity might be better described through a mass-effect perspective, were metacommunity dynamics are mostly determined by dispersal (Leibold et al. 2004).

### Environmental selection

Consistent with previous studies of floral microbiomes, we found that flowers of the yellow monkeyflower (*M. guttatus*) have microbial communities that are distinct from other plant organs (Fig. S6; Junker et al. 2011; Ottesen et al. 2013; Junker & Keller 2015; Wei & Ashman 2018). We provide evidence that floral organs act as different environmental habitats contributing to the assembly of flower microbiomes, despite the small size of *M. guttatus* flowers, and the close contact between stamens and styles with the petals. These results confirm predictions based on knowledge of chemical and morphological differences of these floral parts (Aleklett et al. 2014; Junker & Keller 2015). Specifically, 1) floral organ explains more of the variation in community assembly than seep or pollination treatment, 2) OTUs within a particular flower organ are more phylogenetically clustered than expected by random, and 3) most differences in community composition do not correlate with distance at the scale of this study. Moreover, a neutral model fails to explain the patterns of distribution of a large proportion of OTUs within the flower. Thus, our results corroborate previous work showing that floral organs support different microbial communities (Junker & Keller 2015) and that flower microenvironments (i.e., nectar) can act as strong environmental filters (Herrera et al. 2010) but also extend them by separating the effect of different organs and measuring the effect of dispersal on the effectiveness of organ selection.

We found that a large proportion of OTUs were shared among two or more organs (more than expected by chance for OTUs that deviate from a neutral expectation) and that differences in epiphytic bacterial community composition across organs could be accounted by the nestedeness component of beta diversity. These observations suggest that each organ microbiome is a subset of monkeyflower metacommunity and that, potentially, each organ filter acts in a (more or less) sequential manner. Within the flower, we hypothesize that petals act as the first environmental filter. Petal microbial communities had the highest proportion of unique taxa and showed the strongest signals of selection (i.e., they had a larger proportion of over and underrepresented taxa; Fig. 3B). While some bacteria taxa might be enriched in the flowers, were they are able to grow on floral volatiles and other carbon compounds (Abanda-Nkpwatt et al., 2006), it is likely that the strongest selection is to get rid of potential pathogens and other microbes with potentially negative effects on the plant fitness. Monkeyflower petals have a much larger area than stigmas or stamens and are exposed to a larger proportion of microbes coming from neighboring flowers, transferred by bees, or moved passively from the soil and other organs of the plant. However, once on the petals, microbes could be filtered by petal traits (like pigments, volatiles, trichomes and epidermal cell shapes) that can affect antibiotic properties, surface water retention and temperature of the flower (Whitney et al. 2011; Harrap et al. 2017; Cisowska et al. 2011; Huang et al. 2011). From the petals microbes could colonize the style and stamens. In the case of the style at least, the presence of pollinators increases the rates of microbe colonization from the petals: style communities in the control (open to pollinators) treatment tend to be more similar to petal communities and share more OTUs than those in the pollination exclusion treatment (Fig. 2A; Fig. 2C).

### Dispersal

While it is often assumed that pollinators play a key role in dispersal of microbial communities, and some systems bear this out (e.g. yeast nectar communities; Pozo et al. 2014; Vannette & Fukami 2017), insects visiting flowers have diverse interactions with organs within a flower (Plowright & Laverty 1984). In this study, pollinator treatment on its own did not account for differences in community composition, but rather affected the importance of organ selection in explaining differences in community composition across samples. A similar study looking at community composition of microbiomes of whole tomato flowers found no differences between pollinator exclusion and their control treatments but found increased variation across flower microbiomes in the control (pollinators allowed) relative to flowers in the absence of pollinators (Allard et al. 2018). Here we show that the magnitude of the effect depends on the floral organ as well as the rate and type of interaction with pollinators.

The interplay of dispersal and environmental selection has been hard to disentangle in the field (Evans et al. 2017). Here, we showed that differences in visitation rate of insects to yellow monkeyflower explained some of the variation in treatment differences among locations for the stamen samples. However, the intimacy of the interaction also played a role. Some of these visitors were butterflies and flies that rarely contacted internal organs of the flowers (Fig. 4B), while others had more extensive internal contact with the floral organs. Indeed, the bacterial communities of stamens and styles showed the largest differences between pollination treatments and, in particular, the bacterial communities in the styles had a larger proportion of unique OTUs in the exclusion treatment relative to the control.

*Mimulus guttatus* flowers in the field are mostly visited by medium and large bees foraging for pollen (Robertson et al. 1999; Wu et al. 2008; Arceo-Gómez et al. 2018). Thus, pollinators are likely to have sustained engagement with the stamens and alter the microbial environment of these organs by removing pollen. In a recent paper, Russell et al. (2019) showed that in flowers of *M. guttaus* scrabbling (one of the common behaviors to forage pollen in bees) results in a larger deposition of microbes than other behaviors, and in artificial flowers this behavior leads to the largest deposition of bacteria on the stamen. Here, we showed that differences in bacterial community composition in the stamens across pollinator treatments increased with increased pollination rates. This correlation was clearer when we considered abundance and phylogenetic distance in our analysis. These results are consistent with our observation that, in the absence of pollinators, phylogenetic diversity of the bacterial communities of stamens is much lower than in the presence of pollinators. Finally, the bacterial communities of the stamens in the presence of pollinators are more consistent with a neutral pattern than in our exclusion treatment, and consistent with a contribution of local dispersal, bacterial communities in the anthers became more differentiated with increased distance.

Overall, in this system, petals seem to be acting as a major environmental filter where only a few bacterial taxa can establish and, while a few new bacteria might colonize at high rates of visitation, many of these taxa will remain in low abundance, unable to establish as part of the main petal community. Instead, pollinator engagement with the stamens can introduce variation in the communities and outweigh some of the contributions of environmental selection.

One caveat however, is that amplicon studies can underestimate the effect of organ selection because it is not possible to distinguish dormant species, which can represent a large proportion of microbial communities (Jones and Lennon 2010; Lennon and Jones 2011). Dormancy can facilitate dispersal (Locey 2010) and minimize the experienced environmental stressors (Jones and Lennon 2010) potentially obscuring signals of organ level selection. With the exception of some nectar yeasts and bacteria, and some floral pathogens, we do not know the extent to which microbes are actively growing in flowers. Of the orders we observed in high abundance, many (e.g., Bacillales, Clostridiales, Actinomycetales) are characterized by taxa with spores or other forms of dormancy (e.g., Paredes-Sabja et al. 2011). Future studies should address the proportion of dormant cells in different organs of the flower, the relative contribution of pollinators to that dormant pool as well as the functional roles of microbes in different parts of the flower.

Furthermore, we did not see a signal of community differentiation by distance (except in the stamens of the control treatment), and while one might be tempted to conclude that microbes are “everywhere”, it might instead reflect limited resolution of 16S sequences within the spatial scales chosen in this study. The small sizes of microbes could mean different spatial scales at which environmental selection and dispersal shape local communities. On the one hand, small cells mean that even within a single organ within the plant, microbes might be experiencing many different environments (Lindow & Brandl 2003). The strongest signal of niche sorting might occur at the sub-organ scales. On the other hand, because of their small size dispersal of some microbes might occur at much larger scales than the ones considered in this project (Wilkinson et al. 2012; Choudoir et al. 2018).

Differentiation by distance alone is not the best indicator of neutrality because different patterns of dispersal will result in different patterns of spatial differentiation of communities (e.g. pollinators might travel only a few meters or more than a kilometer in a single dispersal event; Castilla et al. 2017) and locations separated by distance might also experience slightly different environments. This might also be why we did not observe a relationship between the coflowering community at each site and the microbial communities in the flowers of *M. guttatus* in those same sites. In this study we tried to minimize these effects by having the small and medium length distances replicated along different environments and directions (Fig. 1A, B) and by using different lines of evidence to asses neutrality (see *Environmental Selection* section).

While this study provides explicit measurements of neutral and selective contributions of microbial communities of flowers in the presence and absence of pollinators, it also highlights that most of the variation in community composition of floral microbiomes remains unexplained. Factors outside the flower (including soil chemistry) could affect the local pool of microbes or even the floral chemistry (Majetic et al. 2008; Meindl et al. 2014). Similarly, variation across flowers, across plants, or even within a single plant due to competition and strong priority effects (e.g. Peay et al. 2012) could be contributing to the unexplained effects and unfortunately, much of that variation is obscured in this study because to obtain enough DNA we had to pool together the organs of three different flowers from the same cage for each sample.

Here we have shown the importance of “intimacy” and rate of pollination for microbial dispersal in different organs. However, consistent with recent results (Allard et al. 2018), we have also shown that floral communities have a similar composition in the presence or absence of pollinators indicating the importance of other mechanisms of microbial dispersal in shaping floral colonization (e.g., wind, soil and rain). These unknown sources (wind, water, florivores, nectar robbers, other nearby flowers) all could have contributed to the large proportion of OTUs that we were unable to assign to a known source.

Unfortunately, another source of unknown microbes that can play a role (especially for small samples like some of the ones used in this study) is contamination during sampling, extraction and sequencing. While is possible that we had some contamination (it is common in low biomass microbiome analyses; Eisenhofer et al. 2018) we were unable to amplify any of our controls, and to minimize the effects of minor contaminants, we randomized our samples and used sterile equipment at every stage of the process. Similarly, while contamination could explain the presence of some human associated taxa, it could also due to imperfect taxonomy assignment that depends what is already in the database. Additionally, it could be that the species present (which were not able to identify) can be found in flowers. While most of what we know of *Streptoccocus* comes from human pathogens, this genus has also been found in the aerial surfaces of plants (e.g. Pontonino et al., 2018).

Finally, the best practices in analyzing microbiome data are still subject of much debate (e.g. Callahan et al. 2017; Pollock et al. 2018). In this study we analyzed the data in multiple ways (standardizing libraries vs. rarefying data; using OTUs vs. error learning ASV assignment; see methods) and in most cases the method did not affect the results, and in the cases in which it often provided different levels of resolution and different information (same when varying diversity indices). In some cases, these multiple analyses provided added confidence in the results (e.g. community differences between organs). Whereas in other cases (e.g. the relationships between visitation rates and pairwise beta diversity in petal and style samples) discrepancies between analyses suggest caution is warranted.

This study advances our understanding of community assembly of flower microbiomes. It highlights the interplay between dispersal and environmental selection, providing insight into potential effects of pollinator disturbance or floral changes on microbial community composition. As the reproductive structure of angiosperms, microbial effects on flowers can have a large impact on plant fitness. From previous studies we know that microbes of flowers can modify volatile production and nectar composition affecting pollinator visitation (e.g. Herrera et al. 2008; Rering et al. 2017). In addition, the flower is the main site of infection of important pathogens of plants (e.g. anther smut, *Erwinia amylovora*) and microbial communities of flowers can affect the probability of infection (e.g. *Pseudomonas fluorescens* growth on significantly reduces the establishment of *Erwinia*; Wilson and Lindow, 1992).

Despite its importance, the flower remains a relatively understudied environment for microbes and there is still much we do not know. Future studies should address the effects of different floral traits and floral heterogeneity in the assembly of microbial communities, the importance of these microbes to plant fitness and the effects of microbial community assembly on plant communities and the evolution of plant traits. A better understanding of the processes affecting community assembly of flower-associated microbiomes provides insight into the processes driving flower-microbe-pollinator interactions and the potential effects of different disturbances and environmental changes in changing these dynamics.

## Aknowledgments

We are grateful with R. Hayes for field assistance and with P. Aigner and C. Koehler for the facilities provided at the McLaughlin Natural Reserve. Instructors at the EDAMAME workshop, A. Johnson and C. Marshall all provided helpful advice for the analyses. Members of the Ashman lab provided useful feedback at different parts during this project. Feedback from two anonymous reviewers greatly improved this manuscript. This work was supported by the National Science Foundation (DEB 1452386 to T-L. Ashman). M. Rebolleda-Gómez was supported by the Dietrich School of Arts and Sciences through a Pittsburgh Ecology and Evolution Postdoctoral fellowship.

## Data availability

DNA sequences will be made available through NCBI’s Short Read Archive, all other data will be deposited on Dryad. R code for analyses is available on GitHub at github.com/mrebolleda/OrganFilters_MimulusMicrobiome

## Authorship

MRG and TLA designed the research. MRG conducted observations, sampling and analyses. MRG wrote the manuscript with substantial input from TLA.

