## Supplemental figures for "Floral organs act as environmental filters and interact with pollinators to structure the yellow monkeyflower (*Mimulus guttatus*) floral microbiome"

### Supplementary Tables and Figures

Table S1. Sampling locations with coordinates, pollination rates, floral diversity and community samples.

| Seep | Location | Latitude | Longitude | Mean visitation rate-all (visits/hr) | Mean visitation rate-focal sp. (visits/hr) | Floral diversity (Shannon) | Community samples[1] |
|---|---|---|---|---|---|---|---|
| TP9 | 1 | 38.8641 | -122.4278 | 60 | 22 | 0.32 | 2PLST, ZIVE, DEUL, CARU |
| TP9 | 2 | 38.8647 | -122.4272 | 71 | 25 | 0.42 | 3PLST, LOHU, ZIVE |
| TP9 | 3 | 38.8642 | -122.4275 | 34 | 15 | 0.94 | 3PLST, DEUL, CARU |
| TP9 | 4 | 38.8642 | -122.4272 | 52 | 26 | 0.75 | 3PLST, ZIVE, DEUL |
| RH1 | 1 | 38.8592 | -122.4116 | 72 | 55 | 0.75 | MIGU, PLST, DEUL, COSP, ERLA |
| RH1 | 2 | 38.8592 | -122.4108 | 71 | 51 | 0.48 | 2CRMI, ERLA, LOHU, LAMI |
| RH1 | 3 | 38.8592 | -122.4106 | 75 | 48 | 0.62 | 2PLST, 2CRMI, MIGU |
| RHA | 1 | 38.8586 | -122.4094 | 50 | 37 | 1.13 | 2PLST, CRMI, DEUL, ZIVE |
| RHA | 2 | 38.8581 | -122.4097 | 32 | 21 | 0.97 | DEUL, MINU, PLST, LAMI, CRMI |
| RHA | 3 | 38.8581 | -122.4108 | 61 | 32 | 0.51 | 2ALAM, 2MIGU, THMA |
| RHB | 1 | 38.8575 | -122.4072 | 36 | 27 | 1.01 | MINU, ALAM, DEUL, LAMI, ERLA |
| RHB | 2 | 38.8575 | -122.4075 | 42 | 30 | 0.42 | 2MIGU, 2LOHU, GICA |
| RHB | 3 | 38.8564 | -122.4083 | 35 | 6 | 1.05 | 2MINU, MIGU, ERLA, LOHU, |
| BNS | 1 | 38.8612 | -122.3989 | 60 | 34 | 0.45 | MIGU, SIBE, ZIVE, PLST, TRLA |
| BNS | 2 | 38.8612 | -122.3986 | 31 | 15 | 0.41 | ERLA, MIGU, ZIVE, LOHU, DEUL |
| BNS | 3 | 38.8622 | -122.3992 | 48 | 36 | 0.36 | 2MIGU, 2DEUL, PLST |
| BNS | 4 | 38.8619 | -122.3992 | 54 | 10 | 0.82 | 2CRMI, ERLA, DEUL, LOHU |

[1]ALAM-Alium amplectans, CARU-Castilleja rubicundula , COSP- Collinsia sparsiflora, CRMI-Crypantha microstachys, DEUL- Delphinium uliginosum, ERLA- Eriophyllum lanatum, GICA- Gilia capitatum, LAMI- Lagophylla minor, LOHU- Lotus humistratus, MIGU- Mimulus guttatus, MINU- Mimulus nudatus, PLST- Plagiobothrys stipitatus, SIBE- Sisyrinchium bellum, THMA- Thermopsis macrophylla, THMA- Thermopsis macrophylla, TRLA- Triteleia laxa, ZIVE- Zigedenus venenosos

**Table S2.** Table with PERMANOVA summaries using different beta-diversity indices for ASV data (Org- Floral organ; Pol- Pollinator treatment).

| | | **Sorensen** | | | **Bray-Curtis** | | | **Unifrac** | | | **Weighted Unifrac** | | |
|---|---|---|---|---|---|---|---|---|---|---|---|---|---|
| | **Df** | $F$ | $R^2$ | $p$ | $F$ | $R^2$ | $p$ | $F$ | $R^2$ | $p$ | $F$ | $R^2$ | $p$ |
| **Floral organ (Org)** | 2 | 1.45 | 0.03 | 0.001 | 1.643 | 0.034 | 0.001 | 2.973 | 0.06 | 0.001 | 2.389 | 0.048 | 0.001 |
| **Pollinator (Pol)** | 1 | 1.065 | 0.011 | 0.265 | 1.089 | 0.011 | 0.247 | 1.104 | 0.011 | 0.244 | 1.056 | 0.011 | 0.369 |
| **Seep** | 4 | 1.059 | 0.043 | 0.167 | 1.167 | 0.047 | 0.056 | 0.946 | 0.038 | 0.703 | 1.137 | 0.046 | 0.227 |
| **Org*Pol** | 2 | 1.078 | 0.022 | 0.164 | 1.032 | 0.021 | 0.365 | 0.988 | 0.02 | 0.449 | 1.092 | 0.022 | 0.321 |
| **Org*Seep** | 8 | 1.047 | 0.086 | 0.133 | 1.064 | 0.087 | 0.164 | 1.09 | 0.09 | 0.120 | 1.051 | 0.085 | 0.344 |
| **Pol*Seep** | 4 | 1.004 | 0.041 | 0.463 | 0.93 | 0.038 | 0.788 | 0.95 | 0.038 | 0.627 | 0.785 | 0.032 | 0.890 |
| **Residuals** | 70 | | 0.768 | | | 0.763 | | | 0.748 | | | 0.757 | |

**Table S3.** Summary of neutral model fit across floral organs and pollinator treatments.

| Organ | Pollinator treatment | Reads | Samples | Richness | Neutral | | | Binomial | | |
|---|---|---|---|---|---|---|---|---|---|---|
| | | | | | *$R^{2*}$* | AIC | BIC | *$R^{2*}$* | AIC | BIC |
| **Petals** | Control | 1200 | 14 | 251 | 0.062 | -110.332 | -103.281 | 0 | 140.325 | 147.376 |
| | Exclusion | 1200 | 16 | 272 | 0 | -156.764 | -149.552 | 0 | 146.966 | 154.178 |
| **Stamens** | Control | 1200 | 16 | 259 | 0.147 | -214.459 | -207.345 | 0 | 142.698 | 149.812 |
| | Exclusion | 1200 | 15 | 180 | 0.264 | -85.905 | -79.52 | 0 | 98.495 | 104.881 |
| **Style** | Control | 1200 | 17 | 220 | 0.219 | -178.397 | -171.61 | 0 | 125.435 | 132.222 |
| | Exclusion | 1200 | 14 | 165 | 0.249 | -98.889 | -92.677 | 0 | 99.429 | 105.641 |

*Adjusted $R^2$ values that were negative were assigned 0. Negative $R^2$ values indicate that there is more unpredicted variation from the model, than just using the mean.

**Table S4.** Results of Procrustes permutation test to compare co-flowering communities and the microbial communities at each organ and pollinator treatment.

| Organ | Pollinator treatment | *$m^2$* | p-value |
|---|---|---|---|
| **Stamens** | Control | 0.27 | 0.35 |
| | Exclusion | 0.23 | 0.08 |
| **Petals** | Control | 0.20 | 0.18 |
| | Exclusion | 0.29 | 0.54 |
| **Style** | Control | 0.26 | 0.24 |
| | Exclusion | 0.28 | 0.59 |

**Figure S1.** Read distributions A-B. Histograms of samples of a certain number of reads (after QIIME 1.9 pipeline- A; after DADA2 pipeline- B). In green are the samples with eukaryotic reads, in gray are samples after removing eukaryotic sequences. C-D. Rarefaction curves, dashed line is our rarefaction point to include samples of all organs. (C-OTUs obtained after QIIME 1.9 pipeline; D- ASVs obtained after DADA2 pipeline).

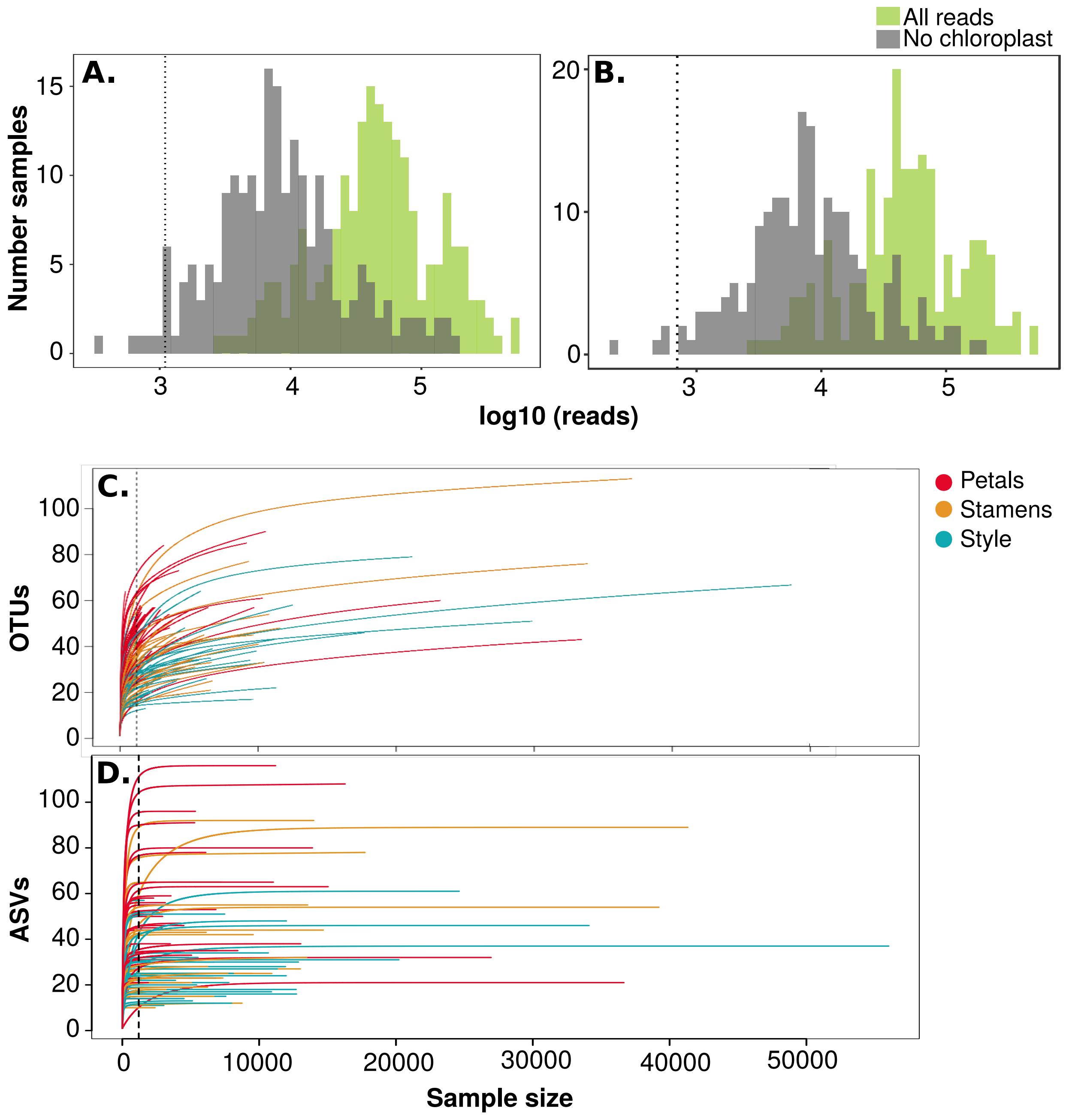

**Figure S2.** Principal coordinate ordination plots showing the contribution of floral organ and seep on microbial community structure. Each point represents one sample, and in different colors are shown the different floral organs sampled. The symbols represent the seep of origin, in solid colors are seeps closer to each other (RH1, RHA, RHB), the distant seeps (TP9, BNS) are in outlined shapes. The left plots (Sørensen and unifrac) are calculated with distances based on presence-absence data alone, the right-side plots (Bray-Curtis and weighted unifrac) are weighted by relative abundances. Similarly, Sørensen and Bray-Curtis take into account only the assigned taxonomy, while unifrac and weighted unifrac include phylogenetic relationships.

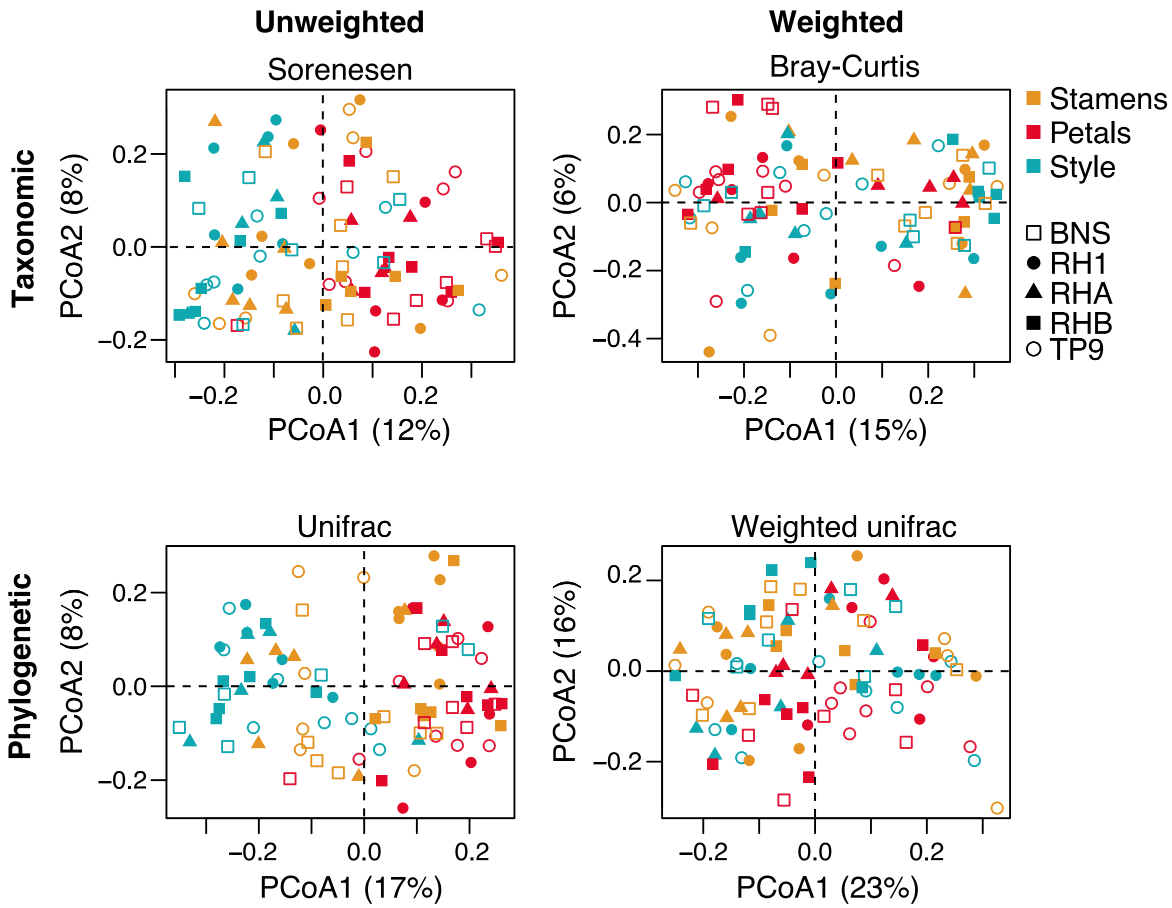

**Figure S3.** Relative abundance of different bacterial orders (those with an overall relative abundance of more than 25%). Each point is one sample, panels on the left are the pollinator exclusion treatments (NP) and panels on the right are samples in the control treatments (WP).

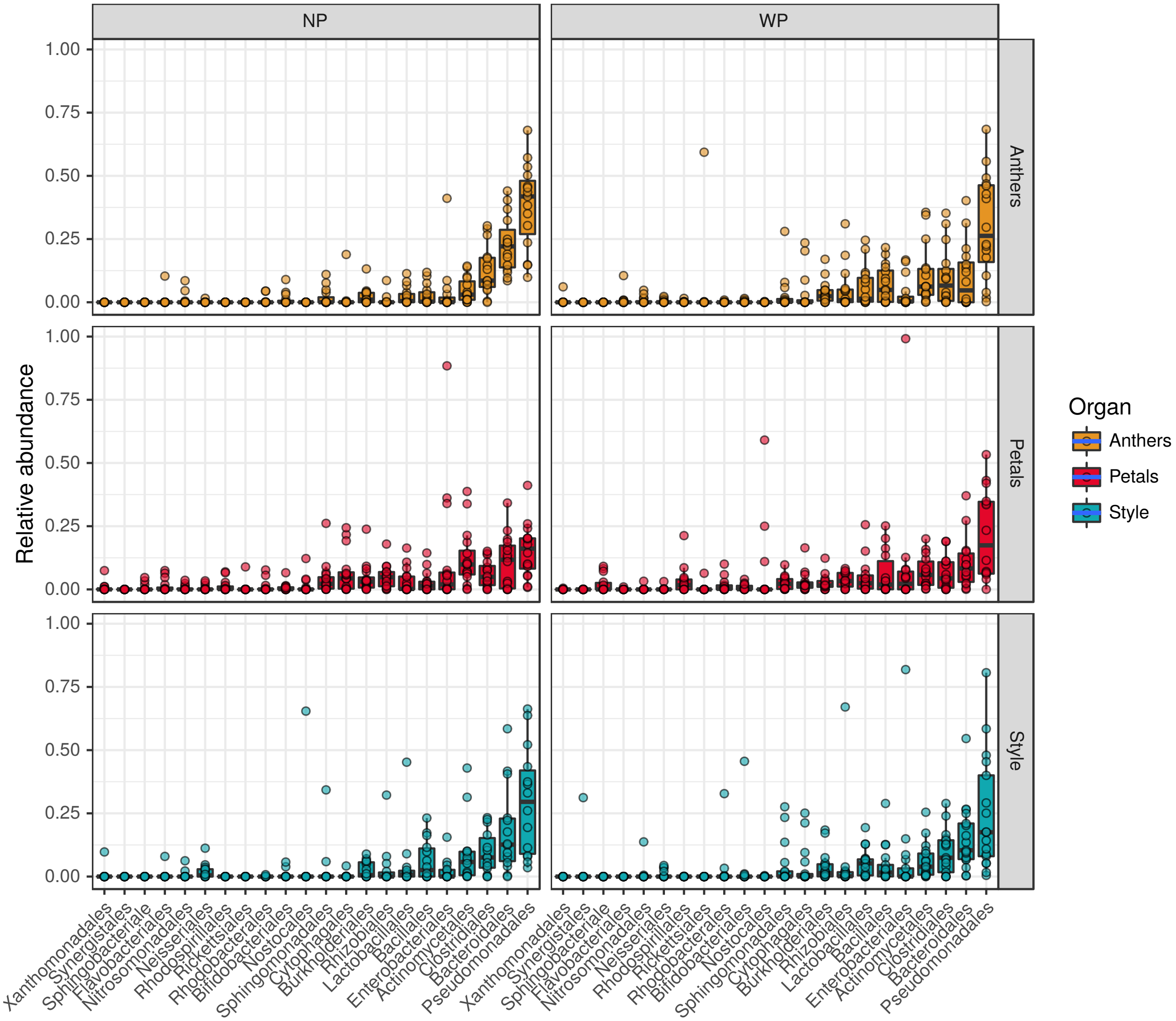

**Figure S4.** Proportion of OTUs shared by at least two organs in each partition (i.e. OTUs distributed under expectations of neutrality, OTUs overrepresented in a particular organ, OTUs underrepresented). The histogram shows the null expectation (given the number of OTUs in each partition) and the dashed line the observed. Upper panels show the data for the pollinator exclusion and the bottom panels the control.

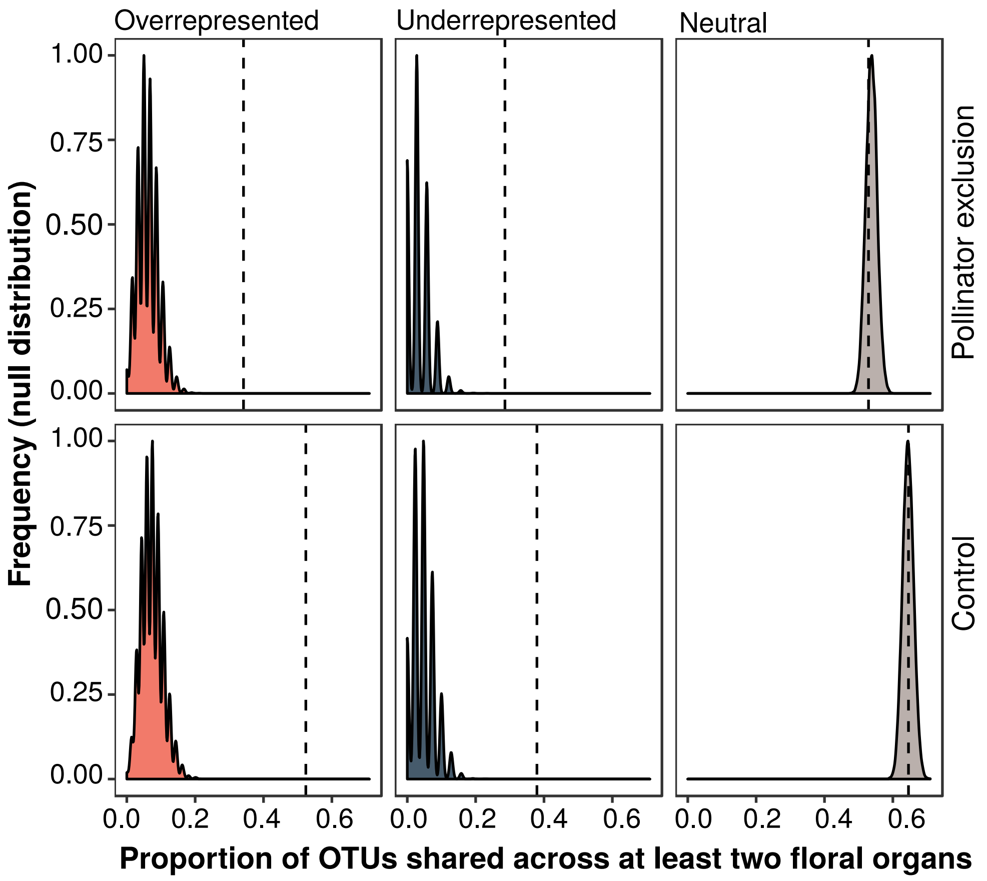

**Figure S5.** Percentage variation in community composition explained by organ in the control and pollinator exclusion (no pollinators) treatments (Sor- Sørensen, B-C- Bray-Curtis, Uni- Unifrac and W-Uni- weighted Unifrac) for the ASV data.

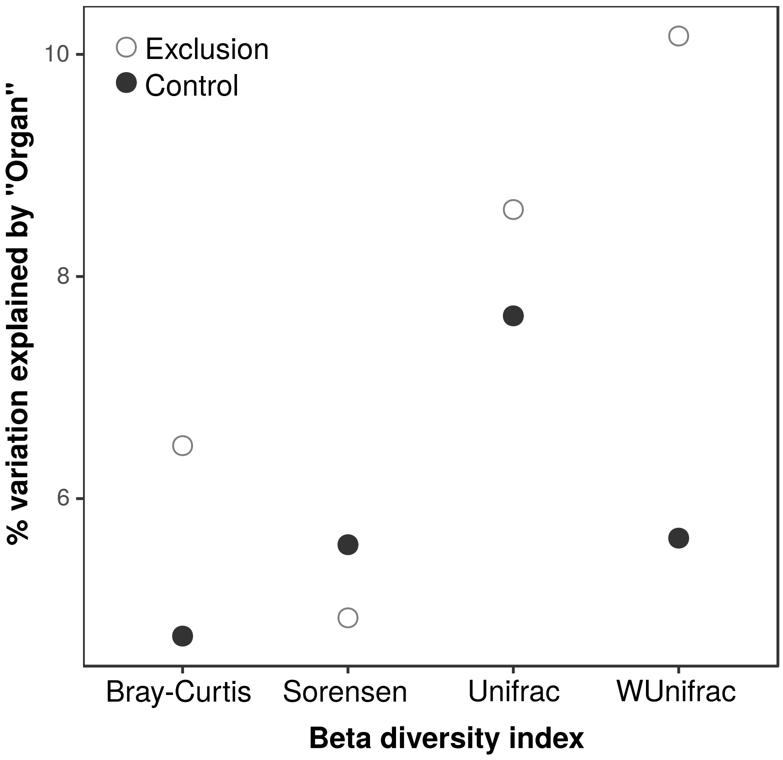

**Figure S6.** The total number of OTUs decays exponentially as we increase the number of samples sharing those OTUs and this decay is the same for the pollinator exclusion treatment and the control. However, overall the exclusion treatment has more OTUs unique to one organ (until the number of OTUs is too small N<20).

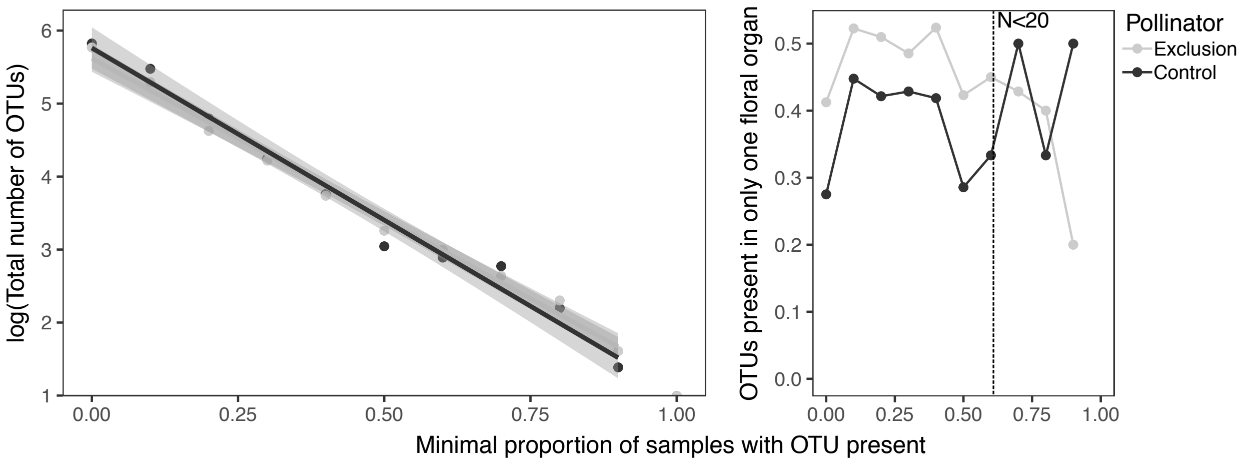


**Figure S7.** Correlations between visitation rate (internal visitors) to the plots and community composition differences between pollinator treatments at each location (pairwise beta diversity). Colors show the strength of the correlation and the numbers inside denote the p-values after adjustment for multiple testing. Analyses were performed using ASV data and different beta diversity indices (Sørensen (Sor) and Unifrac (Uni) include only presence-absence data whereas Bray-Curtis (B-C) and weighted Unifrac (W-Uni) also account for relative abundances).

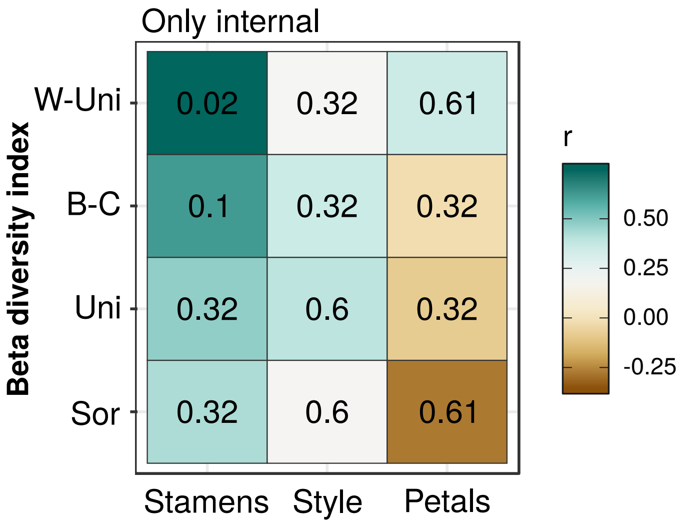

**Figure S8.** Relation of **A.** visitation rate (internal visitors), **B.** co-flowering community diversity and **C.** co-flowering density all with alpha diversity (Shannon index).

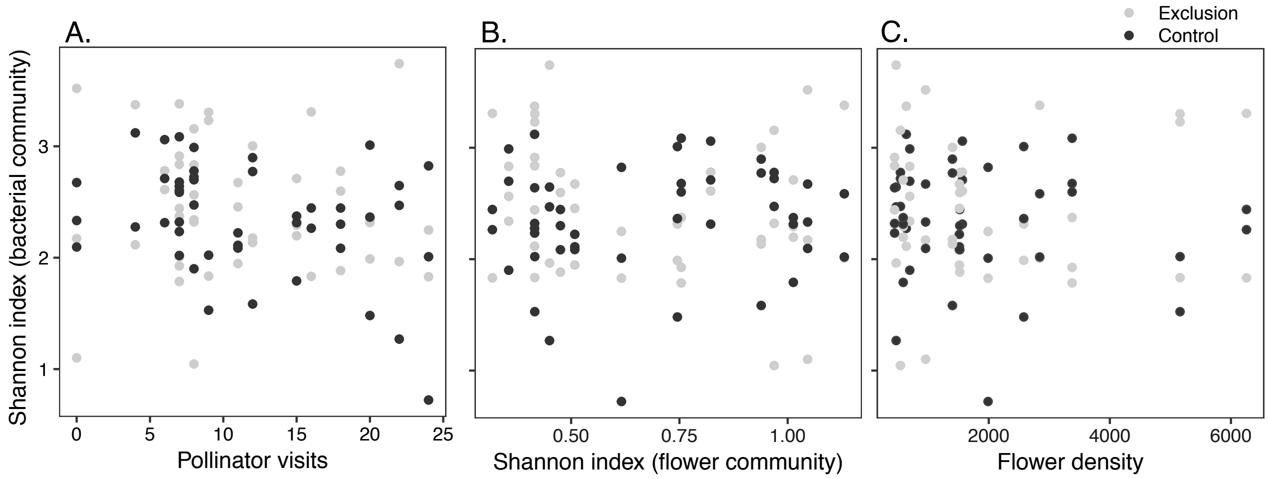
